## Appendix 1 for "Synthesis and Biological Assessment of Chalcone and Pyrazoline Derivatives as Novel Inhibitor for ELF3-MED23 Interaction"

**Method for synthesis of Group 1 to 3**

**General method for synthesis of Group 1 (chalcone analogues)**

A reaction mixture of 3,4,5-trimethoxy acetophenone, benzaldehyde (1.0 equiv.) and 50% NaOH (4.0 equiv.) in EtOH (10mL) was stirred at RT (24 h). 4M HCl (16.0 equiv.) (for 50 and 55) was added to the reaction mixture and kept stirring for 20 min. Water was added and reaction mixture was extracted with ethyl acetate and washed with water and brine, successively, and then dried over anhydrous MgSO_4_. Solvent was removed under reduced pressure and the residue was purified by silica gel column chromatography. (eluent: ethyl acetate:n-hexane)

**(*E*)-3-(4-Hydroxy-3-methoxyphenyl)-1-(3,4,5-trimethoxyphenyl)prop-2-en-1-one (1)**

4-((3,4-Dihydro-2*H*-pyran-6-yl)oxy)-3-methoxybenzaldehyde (1.00 g, 4.27 mmol) and 3,4,5-trimethoxy acetophenone (0.89 g, 4.27 mmol) and 50% NaOH (1.37 mL, 17.08 mmol) were used. Purification was conducted with eluent (ethyl acetate:*n*-hexane = 1:3) to give compound **1** (0.88 g, 59.8%) as an yellow solid. mp 77 - 78 ^o^C; R*_f_* 0.69 (ethyl acetate:*n*-hexane = 1:1); HPLC: R*_T_* 5.39 min (purity; 99.9%); ^1^H-NMR (CDCl_3_, 400 MHz) δ 3.94 (s, 3H), 3.95 (s, 6H), 3.97 (s, 3H), 6.97 (d, *J* = 8.0 Hz, 1H), 7.12 (d, *J* =1.6 Hz, 1H), 7.26 (s, 2H), 7.27 (dd, *J* = 8.0, 1.6 Hz, 1H), 7.31 (d, *J* = 15.2 Hz, 1H), 7.75 (d, *J* = 15.2 Hz, 1H); ^13^C-NMR (CDCl_3_, 100 MHz) 56.3, 56.7, 61.2, 106.4, 110.7, 115.1, 119.8, 123.2, 127.7, 134.1, 142.6, 145.4, 147.0, 148.5, 153.4, 189.7 ppm.

**(*E*)-3-(4-(Allyloxy)-3-methoxyphenyl)-1-(3,4,5-trimethoxyphenyl)prop-2-en-1-one (2)**

4-(Allyloxy)-3-methoxybenzaldehyde (1.03 g, 5.83 mmol) and 3,4,5-trimethoxy acetophenone (1.22 g, 5.83 mmol) and 50% NaOH (1.87 mL, 23.32 mmol) were used. Purification was conducted with eluent (ethyl acetate:*n*-hexane = 1:3) to give compound **2** (1.99 g, 88.4%) as an pale yellow solid. mp 115 - 116 ^o^C; R*_f_* 0.62 (ethyl acetate:*n*-hexane = 1:1); HPLC: R*_T_* 11.30 min (purity; 99.9%); ^1^H-NMR (CDCl_3_, 400 MHz) δ 3.94 (s, 3H), 3.95 (s, 6H), 3.96 (s, 3H), 4.68 (ddd, *J* = 5.2, 1.6, 1.2 Hz, 2H), 5.33 (ddd, *J* = 10.4, 2.4, 1.2 Hz, 1H), 5.43 (ddd, *J* = 17.2, 2.8, 1.6 Hz, 1H), 6.04-6.14 (m, 1H), 6.91 (d, *J* = 8.4 Hz, 1H), 7.16 (d, *J* =2.0 Hz, 1H), 7.23 (dd, *J* = 8.4, 2.0 Hz, 1H), 7.26 (s, 2H), 7.32 (d, *J* = 15.2 Hz, 1H), 7.76 (d, *J* = 15.2 Hz, 1H); ^13^C-NMR (CDCl_3_, 100 MHz) 56.3, 56.7, 61.2, 70.3, 106.4, 111.2, 113.2, 118.7, 120.2, 122.9, 128.3, 132.9, 134.1, 142.6, 145.2, 149.8, 150.7, 153.4, 189.7 ppm.

**(*E*)-3-(4-(But-3-en-1-yloxy)-3-methoxyphenyl)-1-(3,4,5-trimethoxyphenyl)prop-2-en-1-one (3)**

4-(But-3-en-1-yloxy)-3-methoxybenzaldehyde (0.31 g, 1.52 mmol) and 3,4,5-trimethoxy acetophe-none (0.32 g, 1.52 mmol) and 50% NaOH (0.48 mL, 1.92 mmol) were used. Purification was conducted with eluent (ethyl acetate:*n*-hexane = 1:3) to give compound **3** (0.34 g, 54.1%) as an pale yellow solid. mp 133 - 134 ^o^C; R*_f_* 0.69 (ethyl acetate:*n*-hexane = 1:1); HPLC: R*_T_* 13.80 min (purity; 99.9%); ^1^H-NMR (CDCl_3_, 400 MHz) δ 2.50 (dd, *J* = 6.8, 6.8 Hz, 2H), 3.82 (s, 3H), 3.83 (s, 3H), 3.84 (s, 6H), 4.02 (t, *J* = 6.8 Hz, 2H), 5.02 (dd, *J* = 11.2, 2.0 Hz, 1H), 5.08 (dd, *J* = 17.2, 2.0 Hz, 1H), 5.75-5.86 (m, 1H), 6.81 (d, *J* = 8.4 Hz, 1H), 7.07 (d, *J* =2.0 Hz, 1H), 7.14 (dd, *J* = 8.4, 2.0 Hz, 1H), 7.16 (s, 2H), 7.24 (d, *J* = 16.0 Hz, 1H), 7.64 (d, *J* = 16.0 Hz, 1H); ^13^C-NMR (DMSD-*d*_6_, 100 MHz) 33.0, 55.9, 56.2, 60.1, 67.5, 106.2, 112.1, 112.8, 117.1, 119.7, 123.4, 127.6, 133.3, 134.7, 141.8, 144.3, 149.1, 150.5, 152.9, 187.9 ppm.

**(*E*)-3-(4-((*E*)-But-2-en-1-yloxy)-3-methoxyphenyl)-1-(3,4,5-trimethoxyphenyl)prop-2-en-1-one (4)**

(*E*)-4-(But-2-en-1-yloxy)-3-methoxybenzaldehyde (0.46 g, 2.23 mmol) and 3,4,5-trimethoxy acetophe-none (0.47 g, 2.23 mmol) and 50% NaOH (0.71 mL, 8.92 mmol) were used. Purification was conducted with eluent (ethyl acetate:*n*-hexane = 1:3) to give compound **4** (0.79 g, 87.6%) as an yellow solid. m.p. 130 - 131 ^o^C; R*_f_* 0.57 (ethyl acetate:*n*-hexane = 1:1); HPLC: R*_T_* 13.69 min (purity; 99.9%); ^1^H-NMR (CDCl_3_, 400 MHz) δ 1.76 (dd, *J* = 6.4, 1.2 Hz, 3H), 3.94 (s, 6H), 3.95 (s, 6H), 4.59 (d, *J* = 6.0 Hz, 2H), 5.74-5.79 (m, 1H), 5.85-5.92 (m, 1H), 6.92 (d, *J* = 8.4 Hz, 1H), 7.15 (d, *J* = 2.0 Hz, 1H), 7.22 (dd, *J* = 8.0, 2.0 Hz, 1H), 7.26 (s, 2H), 7.32 (d, *J* = 15.6 Hz, 1H), 7.76 (d, *J* = 15.6 Hz, 1H); ^13^C-NMR (DMSD-*d*_6_, 100 MHz) 17.5, 55.7, 56.2, 60.2, 68.6, 106.2, 111.9, 119.7, 123.3, 126.2, 127.5, 130.2, 133.3, 141.8, 144.3, 149.1, 150.2, 152.9, 187.9 ppm.

**(*E*)-3-(3-Methoxy-4-((3-methylbut-2-en-1-yl)oxy)phenyl)-1-(3,4,5-trimethoxyphenyl) prop-2-en-1-one (5)**

3-Methoxy-4-((3-methylbut-2-en-1-yl)oxy)benzaldehyde (0.50 g, 2.27 mmol) and 3,4,5-trimethoxy acetophe-none (0.47 g, 2.27 mmol) and 50% NaOH (0.73 mL, 9.08 mmol) were used. Purification was conducted with eluent (ethyl acetate:*n*-hexane = 1:3) to give compound **5** (0.60 g, 61.2%) as an pale yellow solid. mp 115 - 116 ^o^C; R*_f_* 0.56 (ethyl acetate:n-hexane = 1:1); HPLC: R*_T_* 15.71 min (purity; 99.9%); ^1^H-NMR (CDCl_3_, 400 MHz) δ 1.76 (d, *J* = 0.8 Hz, 3H), 1.79 (d, *J* = 0.8 Hz, 3H), 3.94 (s, 3H), 3.95 (s, 9H), 4.65 (d, *J* = 6.4 Hz, 2H), 5.50-5.54 (m, 1H), 6.91 (d, *J* = 8.4 Hz, 1H), 7.15 (d, *J* = 2.0 Hz, 1H), 7.23 (dd, *J* = 8.8, 2.4 Hz, 1H), 7.26 (s, 2H), 7.32 (d, *J* = 15.6 Hz, 1H), 7.76 (d, *J* = 15.6 Hz, 1H); ^13^C-NMR (CDCl_3_, 100 MHz) 18.5, 26.1, 56.3, 56.7, 61.2, 66.1, 106.4, 110.7, 110.9, 112.9, 115.1, 119.6, 119.9, 123.0, 123.2, 127.9, 138.5, 145.3, 149.9, 153.4, 187.9 ppm.

**(*E*)-3-(3-Hydroxy-4-methoxyphenyl)-1-(3,4,5-trimethoxyphenyl)prop-2-en-1-one (6)**

3-Hydroxy-4-methoxybenzaldehyde (1.0 g, 4.27 mmol) and 3,4,5-trimethoxy acetophenone (0.89 g, 4.27 mmol) and 50% NaOH (1.37 mL, 17.08 mmol) were used. Purification was conducted with eluent (ethyl acetate:*n*-hexane = 1:3) to give compound **6** (1.08 g, 73.5%) as an yellow solid. mp 136 - 137 ^o^C; R*_f_* 0.53 (ethyl acetate:*n*-hexane = 1:1); HPLC: R*_T_* 5.50 min (purity; 99.9%); ^1^H-NMR (CDCl_3_, 400 MHz) δ 3.94 (s, 3H), 3.95 (s, 9H), 5.72 (brs, 1H), 6.89 (d, *J* = 8.4 Hz, 1H), 7.14 (dd, *J* = 8.0, 2.0 Hz, 1H), 7.29 (s, 2H), 7.31 (d, *J* = 2.4 Hz, 1H), 7.35 (d, *J* = 15.6 Hz, 1H), 7.75 (d, *J* = 15.6 Hz, 1H); ^13^C-NMR (CDCl_3_, 100 MHz) 51.1, 56.3, 56.6, 61.2, 106.2, 110.8, 112.9, 120.1, 123.2, 128.8, 133.9, 142.6, 144.9, 146.1, 149.1, 153.4, 189.3 ppm.

**(*E*)-3-(3-(Allyloxy)-4-methoxyphenyl)-1-(3,4,5-trimethoxyphenyl)prop-2-en-1-one (7))**

3-(Allyloxy)-4-methoxybenzaldehyde (0.77 g, 4.00 mmol) and 3,4,5-trimethoxy acetophenone (0.84 g, 4.00 mmol) and 50% NaOH (1.28 mL, 16.00 mmol) were used. Purification was conducted with eluent (ethyl acetate:*n*-hexane = 1:3) to give compound **7** (0.85 g, 55.3%) as an yellow solid. mp 89 - 90 ^o^C; R*_f_* 0.67 (ethyl acetate:*n*-hexane = 1:1); HPLC: R*_T_* 10.85 min (purity; 99.9%); ^1^H-NMR (CDCl_3_, 400 MHz) δ 3.93 (s, 3H), 3.94 (s, 3H), 3.95 (s, 6H), 4.68 (dt, *J* = 5.6, 1.6 Hz, 2H), 5.33 (ddt, *J* = 10.8, 2.8, 1.6 Hz, 1H), 5.45 (ddt, *J* = 16.8, 2.8, 1.6 Hz, 1H), 6.07-6.16 (m, 1H), 6.92 (d, *J* = 8.4 Hz, 1H), 7.18 (d, *J* = 2.0 Hz, 1H), 7.25 (dd, *J* = 8.4, 2.0 Hz, 1H), 7.26 (s, 2H), 7.30 (d, *J* = 15.6 Hz, 1H), 7.75 (d, *J* = 15.6 Hz, 1H); ^13^C-NMR (DMSO-*d*_6_, 100 MHz) 55.7, 56.2, 60.2, 69.1, 106.2, 111.9, 113.2, 117.8, 119.7, 123.7, 127.5, 133.3, 133.8, 141.8, 144.3, 147.7, 151.5, 152.9, 187.9 ppm.

**(*E*)-3-(3-(But-3-en-1-yloxy)-4-methoxyphenyl)-1-(3,4,5-trimethoxyphenyl)prop-2-en-1-one (8)**

3-(But-3-en-1-yloxy)-4-methoxybenzaldehyde (0.10 g, 0.48 mmol) and 3,4,5-trimethoxy aceto-phenone (0.10 g, 0.48 mmol) and 50% NaOH (0.15 mL, 1.92 mmol) were used. Purification was conducted with eluent (ethyl acetate:*n*-hexane = 1:3) to　give compound **8** (0.10 g, 54.2%) as an yellow solid. m.p. 104 - 105 ^o^C; R*_f_* 0.67 (ethyl acetate:*n*-hexane = 1:1); HPLC: R*_T_* 13.13 min (purity; 99.9%); ^1^H-NMR (CDCl_3_, 400 MHz) δ 2.59-2.67 (m, 2H), 3.92 (s, 3H), 3.94 (s, 3H), 3.95 (s, 6H), 4.13 (t, *J* = 6.8 Hz, 2H), 5.14 (ddt, *J* = 10.4, 3.2, 1.2 Hz, 1H), 5.21 (ddt, *J* = 17.2, 3.2, 1.2 Hz, 1H), 5.89-5.99 (m, 1H), 6.92 (d, *J* = 8.8 Hz, 1H), 7.18 (d, *J* = 1.6 Hz, 1H), 7.25 (dd, *J* = 8.8, 1.6 Hz, 1H), 7.29 (s, 2H), 7.30 (d, *J* = 15.6 Hz, 1H), 7.75 (d, *J* = 15.6 Hz, 1H); ^13^C-NMR (DMSD-*d*_6_, 100 MHz) 33.1, 55.7, 56.2, 60.2, 67.7, 106.2, 111.9, 113.1, 116.9, 119.7, 123.5, 127.6, 133.3, 134.9, 141.8, 144.3, 148.1, 151.5, 152.9, 187.9 ppm.

**(*E*)-3-(3-((E)-But-2-en-1-yloxy)-4-methoxyphenyl)-1-(3,4,5-trimethoxyphenyl)prop-2-en-1-one (9)**

(*E*)-3-(But-2-en-1-yloxy)-4-methoxybenzaldehyde (0.50 g, 2.42 mmol) and 3,4,5-trimethoxy aceto-phenone (0.51 g, 2.42 mmol) and 50% NaOH (0.76 mL, 12.10 mmol) were used. Purification was conducted with eluent (ethyl acetate:*n*-hexane = 1:3) to give compound **9** (0.46 g, 47.8%) as an yellow solid. m.p. 92 - 93 ^o^C; R*_f_* 0.57 (ethyl acetate:*n*-hexane = 1:1); HPLC: R*_T_* 13.13 min (purity; 99.9%); ^1^H-NMR (CDCl_3_, 400 MHz) δ 1.77 (dd, *J* = 6.8, 1.6 Hz, 3H), 3.93 (s, 3H), 3.94 (s, 3H), 3.95 (s, 6H), 4.59 (d, *J* = 6.4 Hz, 2H), 5.71-5.83 (m, 1H), 5.86-5.94 (m, 1H), 6.91 (d, *J* = 8.4 Hz, 1H), 7.17 (d, *J* = 2.0 Hz, 1H), 7.24 (dd, *J* = 8.4, 2.0 Hz, 1H), 7.26 (s, 2H), 7.30 (d, *J* = 15.6 Hz, 1H), 7.75 (d, *J* = 15.6 Hz, 1H); ^13^C-NMR (DMSD-*d*_6_, 100 MHz) 17.5, 55.6, 56.2, 60.2, 68.8, 106.2, 111.8, 112.9, 119.6, 123.5, 126.4, 127.4, 130.1, 133.3, 141.8, 144.3, 147.8, 151.5, 152.9, 187.9 ppm.

**(*E*)-3-(4-Methoxy-3-((3-methylbut-2-en-1-yl)oxy)phenyl)-1-(3,4,5-trimethoxyphenyl) prop-2-en-1-one (10)**

4-Methoxy-3-((3-methylbut-2-en-1-yl)oxy)benzaldehyde (0.50 g, 2.27 mmol) and 3,4,5-trimethoxy acetophenone (0.48 g, 2.27 mmol) and 50% NaOH (0.73 mL, 9.08 mmol) were used. Purification was conducted with eluent (ethyl acetate:*n*-hexane = 1:3) to give compound **10** (0.64 g, 47.8%) as an yellow solid. mp 116 - 117 ^o^C; R*_f_* 0.62 (ethyl acetate:*n*-hexane = 1:1); HPLC: R*_T_* 15.05 min (purity; 99.9%); ^1^H-NMR (CDCl_3_, 400 MHz) δ 1.78 (s, 3H), 1.79 (s, 3H), 3.92 (s, 3H), 3.94 (s, 3H), 3.95 (s, 6H), 4.65 (d, *J* = 6.8 Hz, 2H), 5.52-5.56 (m, 1H), 6.90 (d, *J* = 8.4 Hz, 1H), 7.17 (d, *J* = 1.6 Hz, 1H), 7.25 (dd, *J* = 8.4, 1.6 Hz, 1H), 7.26 (s, 2H), 7.31 (d, *J* = 15.6 Hz, 1H), 7.75 (d, *J* = 15.6 Hz, 1H); ^13^C-NMR (DMSD-*d*_6_, 100 MHz) 18.0, 25.4, 55.6, 56.2, 60.2, 65.1, 106.1, 111.7, 112.9, 119.6, 123.5, 127.4, 133.3, 137.3, 141.8, 144.4, 147.9, 151.5, 152.9, 187.9 ppm.

**General method for synthesis of Group 2 (pyrazoline analogues)**

A reaction mixture of chalcone analogue and hydrazine H_2_O(65%) (4.0 equiv.) in AcOH (10 mL) was refluxed (2 h) and then cooled to room temperature. There action mixture was poured into ice and kept overnight at room temperature. Solid formed was filtered and washed with H_2_O. Solid was dried and purified by silica gel column chromatography (eluent: methanol: chloroform).

**1-(5-(4-Hydroxy-3-methoxyphenyl)-3-(3,4,5-trimethoxyphenyl)-4,5-dihydro-1*H*-pyrazol-1-yl)propan-1-one (11)**

Compound **1** (0.10 g, 0.29 mmol) and hydrazine H_2_O (65%) (0.078 mL, 1.16 mmol) were used. Purification was conducted with eluent (methanol:chloroform = 1:100) to give compound **11** (0.078 g, 65.0%) as an yellow solid. mp 178 - 179 ^o^C; R*_f_* 0.48 (methanol : chloroform = 1:10); HPLC: R*_T_* 3.57 min (purity ; 95.5%); ^1^H-NMR (CDCl_3_, 400 MHz) δ 2.42 (s, 3H), 3.15 (dd, *J* = 17.6, 4.4 Hz, 1H), 3.71 (dd, *J* = 17.6, 12.0 Hz, 1H), 3.87 (s, 3H), 3.89 (s, 3H), 3.91 (s, 6H), 5.51 (dd, *J* = 11.6, 4.4 Hz, 1H), 5.55 (s, 1H), 6.72 (dd, *J* = 8.4, 2.0 Hz, 1H), 6.76 (d, *J* = 1.6 Hz, 1H), 6.85 (d, *J* = 8.0 Hz, 1H), 6.96 (s, 2H); ^13^C-NMR (DMSO-d_6_, 100 MHz) 21.7, 42.3, 55.6, 55.9, 59.3, 60.1, 104.1, 110.0, 115.5, 117.3, 126.7, 133.4, 139.4, 145.7, 147.5, 153.0, 154.2, 167.2 ppm.

**1-(5-(4-(Allyloxy)-3-methoxyphenyl)-3-(3,4,5-trimethoxyphenyl)-4,5-dihydro-1*H*-pyrazol-1-yl)ethanone (12)**

Compound **2** (0.50 g, 1.30 mmol) and hydrazine H_2_O (65%) (0.35 mL, 5.20 mmol) were used. Purification was conducted with eluent (methanol:chloroform = 1 : 80) to give compound **12** (0.33 g, 57.4%) as an yellow syrup. R*_f_* 0.58 (methanol:chloroform = 1:9); HPLC: R*_T_* 7.57 min (purity ; 96.0%); ^1^H-NMR (CDCl_3_, 400 MHz) δ 2.73 (s, 3H), 3.44 (dd, *J* = 17.6, 4.4 Hz, 1H), 4.01 (dd, *J* = 17.6, 11.6 Hz, 1H), 4.15 (s, 3H), 4.19 (s, 3H), 4.21 (s, 6H), 4.87 (d, *J* = 5.2 Hz, 2H), 5.56 (dd, *J* = 10.4, 0.8 Hz, 1H), 5.67 (dd, *J* = 17.2, 1.6 Hz, 1H), 5.84 (dd, *J* = 11.6, 4.4 Hz, 1H), 6.31-6.38 (m, 1H), 7.03 (dd, *J* = 8.4, 2.0 Hz, 1H), 7.07 (d, *J* = 2.0 Hz, 1H), 7.11 (d, *J* = 8.4 Hz, 1H), 7.26 (s, 2H); ^13^C-NMR (DMSO-d_6_, 100 MHz) 21.7, 42.3, 55.6, 56.0, 59.2, 60.1, 69.0, 104.1, 109.9, 113.8, 116.9, 117.4, 126.6, 133.9, 135.3, 139.4, 146.7, 149.1, 153.0, 154.2, 167.3 ppm.

**1-(5-(4-(But-3-en-1-yloxy)-3-methoxyphenyl)-3-(3,4,5-trimethoxyphenyl)-4,5-dihydro-1*H*-pyrazol-1-yl)ethanone (13)**

Compound **3** (0.10 g, 0.25 mmol) and hydrazine H_2_O (65%) (0.07 mL, 1.00 mmol) were used. Purification was conducted with eluent (methanol:chloroform = 1:100) to give compound **13** (0.066 g, 60.0%) as an yellow lsyrup. R*_f_* 0.34 (methanol:chloroform = 1:10); HPLC: R*_T_* 9.02 min (purity ; 99.9%); ^1^H-NMR (CDCl_3_, 400 MHz) δ 2.31 (s, 3H), 2.45 (dt, *J* = 6.8, 1.6 Hz, 2H), 3.69 (s, 3H), 3.73 (s, 3H), 3.82 (s, 6H), 3.96 (t, *J* = 6.8 Hz, 2H), 5.05-5.08 (m, 1H), 5.12-5.18 (m, 1H), 5.49 (dd, *J* = 11.6, 4.4 Hz, 1H), 5.82-5.92 (m, 1H), 6.62 (dd, *J* = 8.4, 2.0 Hz, 1H), 6.81 (d, *J* = 2.0 Hz, 1H), 6.89 (d, *J* = 8.4 Hz, 1H), 7.05 (s, 2H); ^13^C-NMR (DMSO-d_6_, 100 MHz) 21.7, 33.1, 42.3, 55.6, 56.0, 59.2, 60.1, 67.7, 104.1, 110.0, 113.6, 116.9, 117.0, 126.6, 134.9, 135.3, 139.4, 147.1, 149.1, 152.9, 154.2, 167.3 ppm.

**(*E*)-1-(5-(4-(But-2-en-1-yloxy)-3-methoxyphenyl)-3-(3,4,5-trimethoxyphenyl)-4,5-dihydro-1*H*-pyrazol-1-yl)ethanone (14)**

Compound **4** (0.50 g, 1.25 mmol) and hydrazine H_2_O (65%) (0.34 mL, 5.00 mmol) were used. Purification was conducted with eluent (methanol:chloroform = 1:80) to give compound **14** (0.29 g, 50.8%) as an yellow syrup. R*_f_* 0.68 (methanol:chloroform = 1:9); HPLC: R*_T_* 9.02 min (purity ; 98.3%); ^1^H-NMR (CDCl_3_, 400 MHz) δ 1.72 (dd, *J* = 6.0, 1.2 Hz, 3H), 2.43 (s, 3H), 3.14 (dd, *J* = 17.6, 4.8 Hz, 1H), 3.71 (dd, *J* = 16.8, 12.0 Hz, 1H), 3.84 (s, 3H), 3.89 (s, 3H), 3.91 (s, 6H), 4.48 (d, *J* = 6.0 Hz, 2H), 5.54 (dd, *J* = 11.6, 4.4 Hz, 1H), 5.69-5.85 (m, 2H), 6.73(dd, *J* = 8.4, 2.0 Hz, 1H), 6.76 (d, *J* = 2.0 Hz, 1H), 6.81 (d, *J* = 8.4 Hz, 1H), 6.96 (s, 2H); ^13^C-NMR (DMSO-d_6_, 100 MHz) 17.5, 21.7, 42.3, 55.5, 55.9, 59.2, 60.1, 68.7, 104.1, 109.8, 113.5, 116.9, 126.5, 126.6, 129.5, 135.1, 139.4, 146.9, 149.1, 152.9, 154.2, 167.3 ppm.

**1-(5-(3-Hydroxy-4-methoxyphenyl)-3-(3,4,5-trimethoxyphenyl)-4,5-dihydro-1*H*-pyrazol-1-yl)ethanone (15)**

Compound **6** (0.10 g, 0.29 mmol) and hydrazine H_2_O (65%) (0.078 mL, 1.16 mmol) were used. Purification was conducted with eluent (methanol:chloroform = 1:80) to give compound **15** (0.096 g, 82.6%) as an white solid. m.p.182 - 183 ^o^C; R*_f_* 0.45 (methanol:chloroform = 1:10); HPLC: R*_T_* 3.77 min (purity ; 98.0%); ^1^H-NMR (CDCl_3_, 400 MHz) δ 2.42 (s, 3H), 3.12 (dd, *J* = 17.6, 4.4 Hz, 1H), 3.70 (dd, *J* = 17.6, 11.6 Hz, 1H), 3.86 (s, 3H), 3.89 (s, 3H), 3.91 (s, 6H), 5.51 (dd, *J* = 11.6, 4.4 Hz, 1H), 5.59 (s, 1H), 6.75-6.80 (m, 3H), 6.96 (s, 2H); ^13^C-NMR (DMSO-d_6_, 100 MHz) 21.7, 42.2, 55.7, 55.9, 59.0, 60.1, 104.1, 110.0, 112.4, 112.5, 126.6, 135.2, 139.4, 146.7, 146.8, 153.0, 154.2, 167.1 ppm.

**1-(5-(3-(Allyloxy)-4-methoxyphenyl)-3-(3,4,5-trimethoxyphenyl)-4,5-dihydro-1*H*-pyrazol-1-yl)ethanone (16)**

Compound **7** (0.42 g, 1.10 mmol) and hydrazine H_2_O (65%) (0.29 mL, 4.40 mmol) were used. Purification was conducted with eluent (methanol:chloroform = 1:80) to give compound **16** (0.33 g, 68.4%) as an yellow syrup. m.p. ^o^C; R*_f_* 0.58 (methanol:chloroform = 1:10); HPLC: RT 7.32 min (purity ; 95.3%); ^1^H-NMR (CDCl_3_, 400 MHz) δ 2.42 (s, 3H), 3.13 (dd, *J* = 17.2, 4.4 Hz, 1H), 3.70 (dd, *J* = 17.6, 12.0 Hz, 1H), 3.83 (s, 3H), 3.89 (s, 3H), 3.92(s, 6H), 4.58 (d, *J* = 5.6 Hz, 2H), 5.25 (ddt, *J* = 10.8, 2.4, 1.6 Hz, 1H), 5.36 (ddt, *J* = 17.2, 3.2, 1.6 Hz, 1H), 5.52 (dd, *J* = 11.6, 4.4 Hz, 1H), 6.00-6.11 (m, 1H), 6.77-6.82 (m, 3H), 6.96 (s, 2H); ^13^C-NMR (CDCl_3_, 100 MHz) 22.2, 42.7, 56.2, 56.5, 59.9, 61.2, 70.3, 104.2, 111.6, 112.1, 118.3, 127.1, 133.6, 134.7, 140.5, 148.5, 149.2, 153.6, 153.9, 168.9 ppm.

**(*E*)-1-(5-(3-(But-2-en-1-yloxy)-4-methoxyphenyl)-3-(3,4,5-trimethoxyphenyl)-4,5-dihydro-1*H*-pyrazol-1-yl)ethanone (17)**

Compound **9** (0.44 g, 1.10 mmol) and hydrazine H_2_O (65%) (0.29 mL, 4.40 mmol) were used. Purification was conducted with eluent (methanol:chloroform = 1:80) to give compound **17** (0.28 g, 56.6%) as an yellow syrup. R*_f_* 0.68 (methanol:chloroform = 1:9); HPLC: R*_T_* 8.29 min (purity ; 99.8%); ^1^H-NMR (CDCl_3_, 400 MHz) δ 1.69 (dd, *J* = 6.0, 1.2 Hz, 3H), 2.43 (s, 3H), 3.13 (dd, *J* = 17.6, 4.4 Hz, 1H), 3.71 (dd, *J* = 17.6, 12.0 Hz, 1H), 3.82 (s, 3H), 3.89 (s, 3H), 3.91 (s, 6H), 4.48 (d, *J* = 6.0 Hz, 2H), 5.53 (dd, *J* = 12.0, 4.4 Hz, 1H), 5.67-5.83 (m, 2H), 6.76-6.82 (m, 3H), 6.96 (s, 2H); ^13^C-NMR (CDCl_3_, 100 MHz) 18.0, 22.2, 42.7, 56.2, 56.5, 60.0, 61.2, 69.9, 104.2, 111.2, 111.9, 118.1, 126.3, 128.6, 131.1, 134.6, 140.5, 148.6, 149.2, 153.6, 153.9, 168.9 ppm.

**General method for synthesis of Group 3 (pyrazoline analogues)**

A reaction mixture of chalcone analogue and hydrazine H_2_O (65%) (4.0 equiv.) in propionic acid (10 mL) was refluxed (2h) and then cooled to room temperature. The reaction mixture was poured into ice and kept overnight at room temperature. Solid formed was filtered and washed with H_2_O. Solid was dried and purified by silica gel column chromatography (eluent: methanol:chloroform).

**1-(5-(4-Hydroxy-3-methoxyphenyl)-3-(3,4,5-trimethoxyphenyl)-4,5-dihydro-1*H*-pyrazol-1-yl)ethanone (18)**

Compound **1** (0.10 g, 0.48 mmol) and hydrazine H_2_O (65%) (0.13 mL, 1.92 mmol) were used. Purification was conducted with eluent (methanol:chloroform = 1:80) to give compound **18** (0.088 g, 75.5%) as an pale yellow solid. mp 180 - 181 ^o^C; R*_f_* 0.44 (methanol:chloroform = 1:10); HPLC: R*_T_* 4.75 min (purity ; 99.9%); ^1^H-NMR (CDCl_3_, 400 MHz) δ 1.20 (t, *J* = 7.2 Hz, 3H), 2.81 (q, *J* = 7.2 Hz, 2H), 3.13 (dd, *J* = 17.6, 4.4 Hz, 1H), 3.70 (dd, *J* = 17.6, 11.6 Hz, 1H), 3.87 (s, 3H), 3.89 (s, 3H), 3.91 (s, 6H), 5.50 (dd, *J* = 12.0, 4.8 Hz, 1H), 5.54 (s, 1H), 6.72 (dd, *J* = 8.4, 2.0 Hz, 1H), 6.75 (d, *J* = 1.6 Hz, 1H), 6.85 (d, *J* = 8.0 Hz, 1H), 6.96 (s, 2H); ^13^C-NMR (DMSO-d_6_, 100 MHz) 9.0, 26.8, 42.1, 55.6, 55.9, 59.3, 60.1, 104.0, 109.9, 115.5, 117.3, 126.8, 133.6, 139.4, 145.7, 147.5, 153.0, 154.0, 170.6 ppm.

**1-(5-(4-(Allyloxy)-3-methoxyphenyl)-3-(3,4,5-trimethoxyphenyl)-4,5-dihydro-1*H*-pyrazol-1-yl)propan-1-one (19)**

Compound **2** (0.20 g, 0.26 mmol) and hydrazine H_2_O (65%) (0.14 mL, 1.04 mmol) were used. Purification was conducted with eluent (methanol:chloroform = 1:80) to give compound **19** (0.094 g, 79.7%) as an yellow syrup. R*_f_* 0.37(methanol:chloroform = 1:10); HPLC: R*_T_* 9.37 min (purity ; 97.6%); ^1^H-NMR (CDCl_3_, 400 MHz) δ 1.19 (t, *J* = 7.6 Hz, 3H), 2.81 (q, *J* = 7.6 Hz, 2H), 3.11 (dd, *J* = 17.6, 4.8 Hz, 1H), 3.68 (dd, *J* = 17.6, 12.0 Hz, 1H), 3.83 (s, 3H), 3.88 (s, 3H), 3.89 (s, 6H), 4.55 (dt, *J* = 5.2, 1.6 Hz, 2H), 5.24 (dq, *J* = 10.4, 1.6 Hz, 1H), 5.35 (dq, *J* = 16.0, 1.6 Hz, 1H), 5.51 (dd, *J* = 11.2, 4.4 Hz, 1H), 5.99-6.07 (m, 1H), 6.71 (dd, *J* = 8.0, 2.0 Hz, 1H), 6.73 (d, *J* = 2.0 Hz, 1H), 6.89 (d, *J* = 8.4 Hz, 1H), 6.94 (s, 2H); ^13^C-NMR (DMSO-d_6_, 100 MHz) 9.0, 26.8, 42.1, 55.5, 55.9, 59.3, 60.1, 69.0, 104.1, 109.8, 113.8, 116.9, 117.4, 126.7, 133.9, 135.4, 139.4, 146.7, 149.1, 153.0, 154.0, 170.6 ppm.

**1-(5-(4-(But-3-en-1-yloxy)-3-methoxyphenyl)-3-(3,4,5-trimethoxyphenyl)-4,5-dihydro-1*H*-pyrazol-1-yl)propan-1-one (20)**

Compound **3** (0.10 g, 0.25 mmol) and hydrazine H_2_O (65%) (0.07 mL, 1.00 mmol) were used. Purification was conducted with eluent (methanol:chloroform = 1:80) to give compound **201** (0.063 g, 53.4%) as an yellow syrup. R*_f_* 0.71 (methanol:chloroform = 1:10); HPLC: R*_T_* 4.36 min (purity ; 96.7%); ^1^H-NMR (DMSO-d_6_, 400 MHz) δ 1.07 (t, *J* = 7.6 Hz, 3H), 2.45 (qt, *J* = 6.8, 1.2 Hz, 2H), 2.51 ~ 2.79 (m, 2H), 3.19 (dd, *J* = 17.6, 4.8 Hz, 1H), 3.69 (s, 3H), 3.73 (s, 3H), 3.82 (s, 6H), 3.79 (dd, *J* = 17.6, 6.8 Hz, 1H), 3.96 (t, *J* = 6.8 Hz, 2H), 5.08 (dq, *J* = 10.0, 1.2 Hz, 1H), 5.15 (dq, *J* = 17.2, 1.6 Hz, 1H), 5.48 (dd, *J* = 12.0, 4.0 Hz, 1H), 5.82-5.92 (m, 1H), 6.62 (dd, *J* = 8.4, 2.0 Hz, 1H), 6.80 (d, *J* = 2.0 Hz, 1H), 6.90 (d, *J* = 8.4 Hz, 1H), 7.04 (s, 2H); ^13^C-NMR (DMSO-d_6_, 100 MHz) 9.0, 26.8, 33.1, 42.1, 55.6, 55.9, 59.3, 60.1, 67.7, 104.1, 109.9, 116.9, 117.0, 126.7, 134.9, 135.4, 139.4, 147.1, 149.1, 152.9, 154.0, 170.6 \ ppm.

**(*E*)-1-(5-(4-(But-2-en-1-yloxy)-3-methoxyphenyl)-3-(3,4,5-trimethoxyphenyl)-4,5-dihydro-1*H*-pyrazol-1-yl)propan-1-one (21)**

Compound **4** (0.10 g, 0.25 mmol) and hydrazine H_2_O (65%) (0.07 mL, 1.00 mmol) were used. Purification was conducted with eluent (methanol:chloroform = 1:100) to give compound **21** (0.042 g, 35.2%) as an yellow semisolid. R*_f_* 0.84 (methanol:chloroform = 1:10); HPLC: R*_T_* 4.54 min (purity ; 95.3%); ^1^H-NMR (DMSO-d_6_, 400 MHz) δ 1.07 (t, *J* = 7.6 Hz, 3H), 1.68 (dd, *J* = 6.8, 1.6 Hz, 3H), 2.68-2.81 (m, 2H), 3.19 (dd, *J* = 18.0, 4.4 Hz, 1H), 3.67 (dd, *J* = 17.6, 12.0 Hz, 1H), 3.69 (s, 3H), 3.72 (s, 3H), 3.76 (dd, *J* = 17.2, 6.0 Hz, 1H), 3.80 (s, 6H), 4.41 (d, *J* = 6.0 Hz, 2H), 5.48 (dd, *J* = 11.6, 4.0 Hz, 1H), 5.63-5.70 (m, 1H), 5.76-5.83 (m, 1H), 6.60 (dd, *J* = 8.0, 2.4 Hz, 1H), 6.78 (d, *J* = 2.0 Hz, 1H), 6.88 (d, *J* = 8.0 Hz, 1H), 7.04 (s, 2H); ^13^C-NMR (DMSO-d_6_, 100 MHz) 9.0, 17.5, 26.8, 42.1, 55.5, 56.0, 59.3, 60.1, 68.7, 104.1, 109.7, 113.5, 116.8, 126.6, 126.7, 135.2, 139.4, 146.8, 149.1, 153.0, 154.0, 170.6 ppm.

**1-(5-(3-Methoxy-4-((3-methylbut-2-en-1-yl)oxy)phenyl)-3-(3,4,5-trimethoxyphenyl)-4,5-dihydro-1*H*-pyrazol-1-yl)propan-1-one (22)**

Compound **5** (0.10 g, 0.24 mmol) and hydrazine H_2_O (65%) (0.07 mL, 1.00 mmol) were used. Purification was conducted with eluent (methanol:chloroform = 1:80) to give compound **22** (0.050 g, 43.0%) as an yellow semisolid. R*_f_* 0.39 (methanol:chloroform = 1:10); HPLC: R*_T_* 3.35 min (purity ; 99.8%); ^1^H-NMR (DMSO-d_6_, 400 MHz) δ 1.07 (t, *J* = 6.8 Hz, 3H), 1.67 (s, 3H), 1.73 (s, 3H), 2.69-2.79 (m, 2H), 3.18 (dd, *J* = 18.0, 4.4 Hz, 1H), 3.69 (s, 3H), 3.73 (s, 3H), 3.68 (dd, *J* = 17.6, 11.6 Hz, 1H), 3.82 (s, 6H), 4.46 (d, *J* = 6.8 Hz, 2H), 5.38-5.42 (m, 1H), 5.48 (dd, *J* = 11.6, 4.4 Hz, 1H), 6.61 (dd, *J* = 8.4, 2.0 Hz, 1H), 6.78 (d, *J* = 2.0 Hz, 1H), 6.88 (d, *J* = 8.0 Hz, 1H), 7.04 (s, 2H); ^13^C-NMR (DMSO-d_6_, 100 MHz) 9.5, 18.4, 25.9, 27.2, 42.5, 55.9, 56.4, 59.7, 60.6, 65.4, 104.5, 110.1, 113.9, 117.3, 120.6, 127.1, 135.6, 137.3, 139.8, 147.4, 149.5, 153.4, 154.2, 171.0 ppm.

**1-(5-(3-Hydroxy-4-methoxyphenyl)-3-(3,4,5-trimethoxyphenyl)-4,5-dihydro-1*H*-pyrazol-1-yl)propan-1-one (23)**

Compound **6** (0.10 g, 0.29 mmol) and hydrazine H_2_O (65%) (0.078 mL, 1.16 mmol) were used. Purification was conducted with eluent (methanol:chloroform = 1:80) to give compound **23** (0.059 g, 48.8%) as an white solid. mp 237 - 238 ^o^C; R*_f_* 0.39 (methanol:chloroform = 1:10); HPLC: R*_T_* 4.69 min (purity ; 99.9%); ^1^H-NMR (CDCl_3_, 400 MHz) δ 1.11 (t, *J* = 7.6 Hz, 3H), 2.74 (q, *J* = 7.6 Hz, 2H), 3.03 (dd, *J* = 17.6, 4.4 Hz, 1H), 3.60 (dd, *J* = 17.6, 11.6 Hz, 1H), 3.77 (s, 3H), 3.80 (s, 3H), 3.83 (s, 6H), 5.40 (dd, *J* = 11.6, 4.4 Hz, 1H), 6.60 (s, 1H), 6.65 (dd, *J* = 8.0, 2.4 Hz, 1H), 6.68 (d, *J* = 2.0 Hz, 1H), 6.71 (d, *J* = 8.0 Hz, 1H), 6.88 (s, 2H); ^13^C-NMR (DMSO-d_6_, 100 MHz) 8.9, 26.7, 42.0, 55.7, 56.0, 59.1, 60.1, 104.0, 112.4, 112.5, 116.1, 126.7, 135.3, 139.4, 146.7, 146.8, 153.0, 154.0, 170.4 ppm.

**1-(5-(3-(Allyloxy)-4-methoxyphenyl)-3-(3,4,5-trimethoxyphenyl)-4,5-dihydro-1*H*-pyrazol-1-yl)propan-1-one (24)**

Compound **7** (0.30 g, 0.78 mmol) and hydrazine H_2_O (65%) (0.21 mL, 3.12 mmol) were used. Purification was conducted with eluent (methanol:chloroform = 1:100) to give compound **24** (0.13 g, 37.1%) as an yellow semisolid. R*_f_* 0.41 (methanol:chloroform = 1:10); HPLC: R*_T_* 3.51 min (purity ; 97.5%); ^1^H-NMR (DMSO-*d_6_*, 400 MHz) δ 1.06 (t, *J* = 7.6 Hz, 3H), 2.82 (dq, *J* = 18.4, 7.2 Hz, 2H), 3.18 (dd, *J* = 17.6, 4.4 Hz, 1H), 3.70 (s, 3H), 3.73 (s, 3H), 3.79 (dd, *J* = 18.0, 6.4 Hz, 1H), 3.82 (s, 6H), 4.50 (dd, *J* = 5.2, 1.2 Hz, 2H), 5.22 (dq, *J* = 10.8, 1.6 Hz, 1H), 5.36 (dq, *J* = 15.6, 1.6 Hz, 1H), 5.47 (dd, *J* = 11.6, 4.4 Hz, 1H), 5.97-6.07 (m, 1H), 6.67 (dd, *J* = 8.8, 1.6 Hz, 1H), 6.79 (d, *J* = 2.0 Hz, 1H), 6.90 (d, *J* = 8.0 Hz, 1H), 7.04 (s, 2H); ^13^C-NMR (DMSO-*d_6_*, 100 MHz) 9.0, 26.8, 42.1, 55.7, 56.0, 59.2, 60.1, 69.0, 104.1, 111.3, 112.3, 117.5, 117.6, 126.7, 133.8, 135.0, 139.4, 147.7, 148.3, 153.0, 154.0, 170.6 ppm.

**(*E*)-1-(5-(3-(But-2-en-1-yloxy)-4-methoxyphenyl)-3-(3,4,5-trimethoxyphenyl)-4,5-dihydro-1*H*-pyrazol-1-yl)propan-1-one (25)**

Compound **9** (1.50 g, 3.76 mmol) and hydrazine H_2_O (65%) (1.01 mL, 15.04 mmol) were used. Purification was conducted with eluent (methanol:chloroform = 1:80) to give compound **6** (0.078 g, 4.9%) as an yellow semisolid. R*_f_* 0.39 (methanol:chloroform = 1:9); HPLC: R*_T_* 4.15 min (purity ; 97.5%); ^1^H-NMR (DMSO-*d_6_*, 400 MHz) δ 1.07 (t, *J* = 7.6 Hz, 3H), 1.65 (dd, *J* = 6.4, 1.2 Hz, 3H), 2.75 (dq, *J* = 16.0, 8.0 Hz, 2H), 3.19 (dd, *J* = 18.0, 4.4 Hz, 1H), 3.69 (s, 3H), 3.72 (s, 3H), 3.78 (dd, *J* = 18.0, 6.0 Hz, 1H), 3.82 (s, 6H), 4.41 (d, *J* = 6.0 Hz, 2H), 5.47 (dd, *J* = 11.6, 4.4 Hz, 1H), 5.60-5.68 (m, 1H), 5.73-5.82 (m, 1H), 6.66 (dd, *J* = 8.0, 1.6 Hz, 1H), 6.78 (d, *J* = 2.0 Hz, 1H), 6.88 (d, *J* = 8.4 Hz, 1H), 7.04 (s, 2H); ^13^C-NMR (DMSO-*d_6_*, 100 MHz) 9.0, 17.4, 26.8, 42.0, 55.6, 56.0, 59.2, 60.1, 68.6, 104.1, 110.9, 112.1, 117.3, 126.5, 126.7, 129.8, 134.9, 139.4, 147.8, 148.2, 153.0, 154.0, 170.5 ppm.

**1-(5-(4-Methoxy-3-((3-methylbut-2-en-1-yl)oxy)phenyl)-3-(3,4,5-trimethoxyphenyl)-4,5-dihydro-1*H*-pyrazol-1-yl)propan-1-one (26)**

Compound **10** (0.20 g, 0.48 mmol) and hydrazine H_2_O (65%) (0.13 mL, 1.92 mmol) were used. Purification was conducted with eluent (methanol:chloroform = 1:80) to give compound **26** (0.049 g, 21.3%) as an yellow semi-solid. R*_f_* 0.69 (methanol:chloroform = 1:9); HPLC: R*_T_* 4.62 min (purity ; 95.4%); ^1^H-NMR (DMSO-*d_6_*, 400 MHz) δ 1.07 (t, *J* = 7.6 Hz, 3H), 1.65 (s, 3H), 1.67 (s, 3H), 2.74 (dq, *J* = 16.0, 8.0 Hz, 2H), 3.17 (dd, *J* = 18.0, 4.4 Hz, 1H), 3.69 (s, 3H), 3.71 (s, 3H), 3.75 (dd, *J* = 18.0, 8.8 Hz, 1H), 3.80 (s, 6H), 4.48 (d, *J* = 6.8 Hz, 2H), 5.34 (dd, *J* = 6.4, 5.2 Hz, 1H), 5.47 (dd, *J* = 11.6, 4.0 Hz, 1H), 6.66 (dd, *J* = 8.4, 2.0 Hz, 1H), 6.73 (d, J = 2.0 Hz, 1H), 6.88 (d, *J* = 8.4 Hz, 1H), 7.04 (s, 2H); ^13^C-NMR (DMSO-d_6_, 100 MHz) 9.0, 17.9, 25.3, 26.8, 42.0, 55.6, 56.0, 59.3, 60.1, 64.9, 104.1, 110.7, 112.0, 117.3, 120.1, 126.7, 135.0, 136.8, 139.4, 147.9, 148.3, 153.0, 154.0, 170.5 ppm.
